# Anthropogenic disturbance impacts gut microbiome homeostasis in a Malagasy primate

**DOI:** 10.1101/2022.04.02.486803

**Authors:** Wasimuddin, Hina Malik, Yedidya R Ratovonamana, S Jacques Rakotondranary, Jörg U. Ganzhorn, Simone Sommer

## Abstract

Increasing anthropogenic disturbances in Madagascar are exerting constrains on endemic Malagasy lemurs and their habitats, with possible effects on their health and survival. An important component of health is the gut microbiome, which might be disrupted by various stressors associated with environmental change. We have studied the gut microbiome of grey-brown mouse lemurs *(Microcebus griseorufus)*, one of the smallest Malagasy primates and an important model of the convergent evolution of diseases. We sampled two sites: one situated in a national park and the other consisting of a more disturbed site around human settlement. We found that more intense anthropogenic disturbances indeed disrupted the gut microbiome of this lemur species with a marked reduction in its diversity and led to a shift in microbial community composition. Interestingly, we noted a decrease in beneficial bacteria together with a slight increase in disease-associated bacteria and alterations in microbial metabolic functions. Because of the crucial services provided by the microbiome to host health and in disease, such negative alterations in the gut microbiome of the lemur from anthropogenically disturbed habitat might impact the health of mouse lemurs, rendering them susceptible to diseases and ultimately affecting their survival in the shrinking biodiversity seen in Madagascar. Gut microbiome analyses might thus serve as an early warning signal for pending threats to lemur populations.

## Introduction

Anthropogenic disturbances frequently have direct negative influences on wildlife distribution, reproduction, physiology and health. However, they can also exert indirect effects by changing the abundance pattern of species communities and their interactions, ultimately influencing the population dynamics of wild-living species (Otto, 2018). Often, these disturbances act in synergy, exacerbating the decline of animal populations. Madagascar has been identified as one of the world’s biodiversity hotspots that continues to be under severe anthropogenic pressure (Myers et al., 2000; Vieilledent et al., 2018). High forest fragmentation, encroachment into wildlife areas, habitat degradation and heavy pressures on forest resources are critical problems in Madagascar, because of the exponential rise in its human population coupled with extreme poverty (Morelli et al. 2019; Estrada et al. 2018; World Bank 2021). Rapid modifications of their habitats place immediate pressures on animal populations, with the possible extinction of species that cannot tolerate such alterations (Otto, 2018). Primates are especially susceptible to such disturbances, as primate habitat ranges often overlap extensively with the growing human population. Indeed, out of the 107 lemur species assessed by the IUCN in Madagascar, 103 were considered to be threatened, with 33 of them being listed as critically endangered, making lemurs the most threatened vertebrate taxon in Madagascar (IUCN, 2021). The drivers causing the decline of lemurs can act synergistically, although habitat loss because of cultivation and timber harvesting is the major contributor (Irwine et al. 2011; Schwitzer et al. 2014), which is amplified by the negative impact of anthropogenic disturbance on the physiology and health of primates (Junge et al., 2011; Schwitzer et al., 2011; Bublitz et al., 2015). The response of species to habitat disturbance has shown to be mixed and species-trait dependent, e.g. small-bodied and mixed folivorous/frugivorous primates show a higher resilience compared with large bodied species and those with a more specialized diet (Eppley et al., 2020). However, more resilient species might also suffer from human encroachment, especially if this disturbs the host-gut microbiome (Fackelmann et al. 2021).

Now known as an important component of health, the gut microbiome provides several necessary services to its host, ranging from nutrition uptake and metabolism to the shaping of immunity, disease resistance and behaviour and its overall support of the host’s adaptation to various environment conditions (Thomas et al., 2017; McKenney et al., 2018; Fackelmann et al., 2021; Henry et al., 2021). The gut microbiome itself is shaped by several intrinsic (i.e. genetics) and extrinsic (i.e. environment, social interactions) factors (Amato, 2013; Bahrndorff et al., 2016). Anthropogenic disturbance can induce negative changes in natural gut microbiome homeostasis, because of the loss of microbial diversity, especially a decrease in commensal bacteria and an increase of pathogens, and can result in a state of ‘dysbioses’, with negative implications for animal health. Indeed, recent studies have emphasized the negative effects of landscape modifications and habitat fragmentation on the homeostasis of the host-gut microbiome. For example, the gut microbiome of howler monkeys living in fragmented areas or in captivity has been shown to have a lowered diversity (Amato et al., 2013), with several follow-up studies confirming these findings in various settings and for diverse non-human primate species (Barelli et al. 2015, 2020; Hayakawa et al. 2018; Trosvik et al. 2018). Habitat fragmentation is additionally associated with dietary shifts and microbiota variability in common vampire bats (Ingala et al., 2019). Recent wildlife studies have highlighted that gut homeostasis is also subject to disturbance by viral infections, which facilitate co-infections and the spread of zoonotic diseases (Wasimuddin et al., 2018, 2019), and is influenced by the host’s immune genetic diversity (Montero et al., 2021). The associated behavioural and physiological traits of host species also play an important role in rendering resistance or susceptibility within the gut microbiome (Barelli et al., 2020).

Here, we have studied the impact of anthropogenic disturbances on the gut microbiome of the grey-brown mouse lemur (*Microcebus griseorufus*) from Southern Madagascar. Grey-brown mouse lemurs are small-bodied (40–80 g) arboreal nocturnal primates inhabiting both natural and degraded forests, exhibiting a potential range overlap with humans and other domestic species (Mittermeier et al. 2010). *Microcebus* spp. represent important model species for understanding the convergent evolution of diseases (Ezran et al., 2017; Hozer et al., 2019) and provide a good study system for disentangling the effects of anthropogenic disturbances on the homeostasis of the gut microbiome and on overall host health. The diet of the grey-brown mouse lemur consists of mostly gum and fruits, but also includes insects. In the dry season when the food supply is short, the lemur reduces its metabolic activities and enters torpor or a hibernation-like stage (Mittermeier et al. 2010). Small-bodied primates and nocturnal primates have been suggested to be less sensitive to anthropogenic disturbances (Eppley et al., 2020; Hending, 2021). Nevertheless, anthropogenically modified habitats with substantial contact zones between lemurs and humans increase disturbance frequencies and modify habitat and microclimatic characteristics, including food resources. In order to understand the influence of anthropogenic disturbances on gut microbiome community structure and function, we have compared the gut microbiome of mouse lemurs from two sites: a less disturbed site located in the Tsimanampetsotsa National Park and a highly disturbed site partially integrated into a village with associated livestock herding and agriculture.

## Material and Methods

### Study sites and sample collection

The field study was carried out between 2013 and 2015 in south-western Madagascar at two localities: Andranovao and Miarintsoa (Figure 1). Andranovao is at the western foot of the Mahafaly plateau, located in the Tsimanampetsotsa National Park, in south-western Madagascar (24°03′–24°12′S, 43°46′–43°50′E). The vegetation consists of continuous dry spiny forest and spiny thicket (Yedidya Ratovonamana et al., 2011, 2013). Here, annual rainfall averages 300 mm, with some years being without rain (Yedidya Ratovonamana et al., 2013). Miarintsoa is located on the limestone plateau about 42 km east of Andranovao at 23°50′ S, / 44°06′ E; 183 m a.s.l. The area around Miarintsoa was largely covered with forest until 1973 but has been heavily degraded to a highly fragmented state since 2010 (Brinkmann et al., 2014); it is now dominated by agricultural fields and pasture with small and scattered forest remnants used intensively for wood collection. The area around Miarintsoa can be described as being land covered with grasses (mainly *Heteropogon contortus*), with woody plant cover of 10 - 30% (according to the vegetation classification by Brinkmann et al. 2014; Nopper et al. 2018). On the plateau, annual rainfall ranges between 500 and 800 mm (Yedidya Ratovonamana et al., 2013).

**Figure 1.**
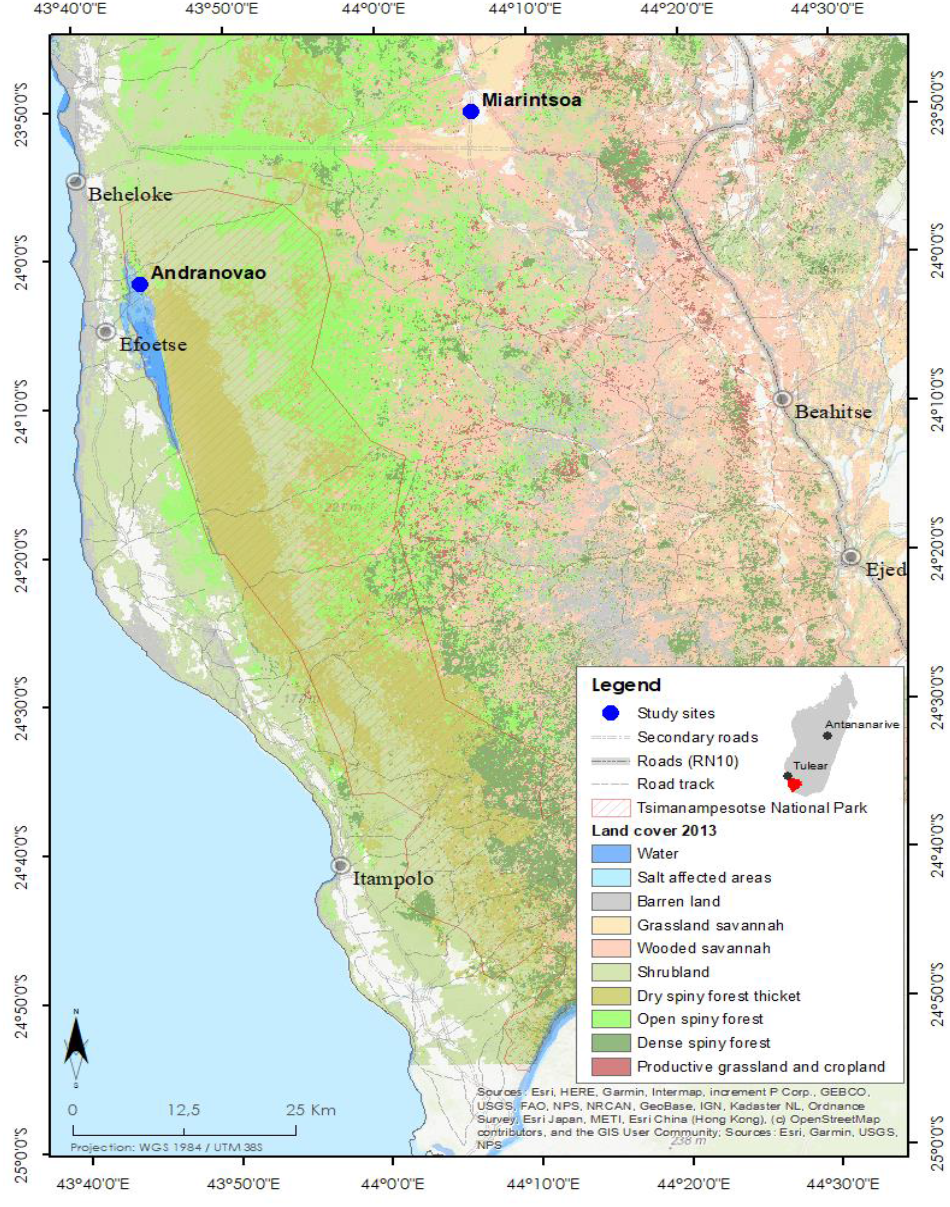
Study area and vegetation cover in south-western Madagascar. The study sites Andranovao and Miarintsoa are marked with blue dots. The Tsimanampetsotsa National Park boundaries and the type of land cover (in 2013) is indicated (adapted from www.sulama.de by Y. R. Ratovonamana).

We used samples from 161 *Microcebus griseorufus* individuals captured with Sherman live-traps (H.B. Sherman Traps, Tallahassee; 7.5 × 7.5 × 30.5 cm) following standard protocols (e.g., Bohr et al. 2011). All trapped animals were sexed, measured, weighed and marked permanently with subdermal transponders. None of the individuals exhibited any signs of illness or disease at the time of capture. Fresh faecal samples were collected either from the traps or handling bags and preserved in Eppendorf vials filled with 500 μl RNAlater (Life Technologies). After morphological measurement, the animals were released immediately at the site of capture. Upon return to the field station, samples were kept cool (around 20 °C) until they could be transported to the nearest city. They were then stored at −20 °C before being transported in polystyrene isolation boxes equipped with passive cooling elements to Germany where they were kept at −80 °C until DNA extraction.

### Bacterial DNA extraction and 16S rRNA gene amplicon sequencing

Faecal samples were homogenized by using beads in a SpeedMill PLUS Homogenizer (Analytik Jena, Germany) (Wasimuddin et al. 2019, Montero et al. 2021). Bacterial genomic DNA was extracted from faeces by using NucleoSpin 96 Soil kits (Macherey-Nagel, Germany) according to the manufacturer’s instructions. We amplified the hypervariable V4 region of the 16S rRNA gene (291 bp) with the primer pair 515F (5-GTGCCAGCMGCCGCGGTAA-3) and 806R (5-GGACTACHVGGGTWTCTAAT-3) (Kuczynski et al., 2011; Caporaso et al., 2012; Wasimuddin et al., 2020). Following the approach recommended for the Fluidigm System (Access Array™ System for Illumina Sequencing Systems, © Fluidigm Corporation), these primers were tagged with sequences (CS1 forward tag and CS2 reverse tag) complementary to the respective forward or reverse access array barcode primers required for the Illumina platform. The polymerase chain reaction (PCR) and PCR-product purification were performed as previously reported (Wasimuddin et al., 2019; Montero et al., 2021). After quantification by DropSense (Trinean, US), samples were pooled to give an equal amount of 50 ng DNA and the pool was diluted to 8 pM in hybridization buffer. The final library was paired-end sequenced in a single run on an Illumina^®^ MiSeq platform. Negative controls were included for both the DNA extraction (no sample added) and 16S PCR amplification (with PCR certified water) to test for contamination. No DNA contamination was observed when using DropSense quantification.

### Bioinformatics

Demultiplexed reads without barcodes and adapters were received as output from the Illumina sequencing platform. All subsequent analyses were performed within the R environment (R version 3.5.1, R Development Core Team 2011). For data pre-processing, we followed the *DADA2* pipeline (version 1.10) (Callahan et al., 2016). For the dataset, reads were trimmed from both ends based on the quality profile; error rates were obtained from the data by using the parametric error model as implemented in *DADA2*. After the reads had been denoised and merged, chimeric sequences were removed from the dataset by following the ‘consensus’ method. Thus, non-chimeric amplicon sequence variants (ASVs; i.e. sequences differing by as little as one nucleotide) were identified in each sample. The taxonomy of representative ASVs was assigned using the naïve Bayesian classifier method with the SILVA v132 non-redundant (NR) database. Species level was designated by the exact matching (100% identity) of ASVs with database sequences, as previously recommended (Edgar, 2018). The *Phyloseq* (version 1.24.2) (McMurdie and Holmes, 2013) package was used for further data processing, with ASVs belonging to chloroplast, mitochondria, archaea, Eukaryota and unassigned ASVs at the phylum level being removed from the dataset. As *DADA2* might be more sensitive to a low amount of contamination (Caruso et al., 2019), we further removed the ASVs showing less than ten reads in the overall dataset. We recovered, on average, 52,136 (range 12,081-130,215) high quality reads per sample. In total, 1503 unique ASVs were detected in the gut microbiomes of grey-brown mouse lemurs (N= 161 individuals) after taxonomic assignments. Since we aimed for a minimum depth of 20,000 reads in analyses requiring rarefaction (alpha & beta diversity calculations), we removed the sample with 12,081 reads from these analyses.

### Alpha and beta diversity analyses and identification of phyla and ASVs associated with anthropogenic disturbance

We investigated the effects of habitat differing in anthropogenic disturbance and of sex on microbial diversity for each sample by using three different alpha diversity indices (number of observed species, Fisher and Shannon) after rarefying the data to the 23,150 sequences per sample (one sample with 12,081 reads was removed before rarefaction) by using *Phyloseq*. The first alpha diversity index describes the unique taxa count (i.e. ASV, species richness), whereas the Fisher index accounts for the number of species and for abundance and the Shannon index is an abundance-weighted diversity measure. To analyse the association of study site and sex with these alpha diversity matrices, we performed General Linear Modelling (GLM) by using the *lme*_*4*_ package in R (Bates et al., 2015). We included the study sites (Andranovao; n= 113, Miarintsoa; n= 47) and sex (male; n= 69, female; n= 89, not known; n= 2) in the model as explanatory variables for each alpha diversity metric table.

Beta diversity analyses were based on calculated Euclidean, Unweighted and Weighted UniFrac dissimilarity matrices after the data had been rarefied to 23,150 sequences per sample by using *Phyloseq*. The Euclidean distance is commonly used to detect changes in relative composition without using phylogenetic information, whereas Unweighted UniFrac distances account for phylogenetic distances among taxa present and Weighted UniFrac distances additionally include information on taxa abundance. The permutational multivariate analysis of variance (PERMANOVA) was employed as implemented in the *adonis* function of the *vegan* package (version 2.5-2) (Oksanen et al., 2018) in order to test the significance of the differences in community composition with 999 permutations. For all beta diversity metrics, we similarly included study site and sex in the models as explanatory variables. To visualize patterns of separation between the different sample categories, principal coordinates analyses (PCoA) were performed based on the Euclidean and Unweighted and Weighted UniFrac dissimilarity matrices.

In order to identify the phyla and ASVs accountable for differences in the microbiome of lemurs inhabiting different habitats, we employed a negative binomial model-based approach available in the *DESeq2* package in R (Love et al., 2014) without rarefaction. Wald tests were performed and only phyla and ASVs remaining significant (p<0.01) after the Benjamini–Hochberg correction were retained. Additionally, the *Random forest* classifier was used to assign the rank of differentially abundant phyla to the microbial community based on the mean decrease in accuracy (Ssekagiri et al., 2017).

### Microbial function predictions and identification of differentially abundant functional pathways

To anticipate functional differences of the gut microbiome of individuals inhabiting the different habitats based on 16S rRNA gene sequencing, we used *PICRUSt2* (Douglas et al., 2019). After normalization for copy number variation as implemented in *PICRUSt2*, the metagenome prediction was carried out by using the KEGG Orthology (KOs) classification. To assess the accuracy of the prediction, weighted Nearest Sequenced Taxon Index (weighted NSTI) scores were calculated. The NSTI score represents the sum of phylogenetic distances of each organism in a sample to its nearest relative in a sequenced bacterial genome. Only sequences with NSTI values of <2, as default, were kept in order to allow better predictions of metagenomes. To investigate the effect of the anthropogenic disturbances on the KEGG composition, we calculated ‘*gower’* distances after rarefying the data. We applied PERMANOVAs with 999 permutations and included study site and sex of the individual in the model as explanatory variables to explain differences in the *gower* metrics. PCoA plots were used to visualize the pattern of separation of metagenomes based on the *gower* metric between individuals captured in Andranovao or Miarintsoa. We categorized KOs to major functional pathways by applying the KEGG classification at hierarchy level 3. To identify the pathways that showed differential abundance between Andranovao and Miarintsoa individuals, we used Wald tests implemented in the *DESeq2* package in R (Love et al., 2014) and only pathways that remained significant (p≤0.05) after the Benjamini–Hochberg correction were reported.

## Results

### Gut microbial diversity and community composition of grey-brown mouse lemurs differ between habitats

The gut bacterial communities of grey-brown mouse lemurs were dominated by the phyla Bacteroidetes (32.3%), Actinobacteria (30.2%) and Firmicutes (25.4%), followed by Proteobacteria (5.9%) and Epsilonbacteraeota (4.3%). We used General Linear Modelling (GLM) to test whether inter-individual differences within three alpha diversity estimates (number of observed species, Fisher and Shannon) can be explained by host habitat and sex. Habitat had a significant effect on the number of observed species (p=0.003), on Fisher (p=0.003) and on Shannon (p=0.016), but not sex (all p>0.05). All three alpha diversity metrics were lower in individuals from Miarintsoa than Andranovao (Figure 2, Supplementary Table S1).

**Figure 2.**
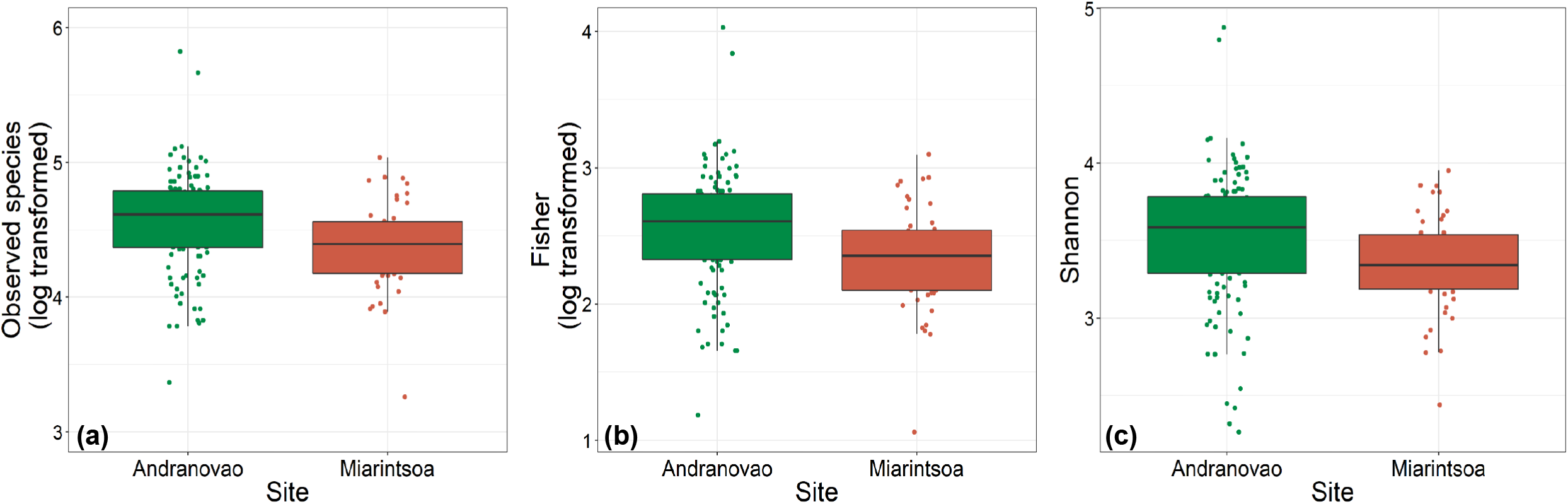
Alpha diversity of gut bacteria in grey-brown mouse lemurs differs between habitats. Effect of study site on (a) number of observed species, (b) Fisher diversity index and (c) Shannon diversity index of mouse lemur individuals. Individuals from Miarintsoa (red) revealed a lower bacterial diversity than individuals trapped in Andranovao (green) (see Supplementary Table 1 for details).

To determine whether the host habitat influenced the composition of the gut microbial community, we calculated three beta diversity metrics (Euclidean, unweighted UniFrac and weighted UniFrac) and included again habitat and sex as explanatory variables in PERMANOVA models. We observed a significant effect of habitat (unweighted UniFrac: R^2^=0.046, p=0.001; weighted UniFrac: R^2^=0.033, p=0.001; Euclidean: R^2^=0.035, p=0.001) but no effect of host sex (unweighted UniFrac: R^2^=0.006, p=0.280; weighted UniFrac: R^2^=0.006, p=0.364; Euclidean: R^2^=0.006, p=0.426) on gut microbial beta diversity estimates (Figure 3, Supplementary Table S2).

**Figure 3.**
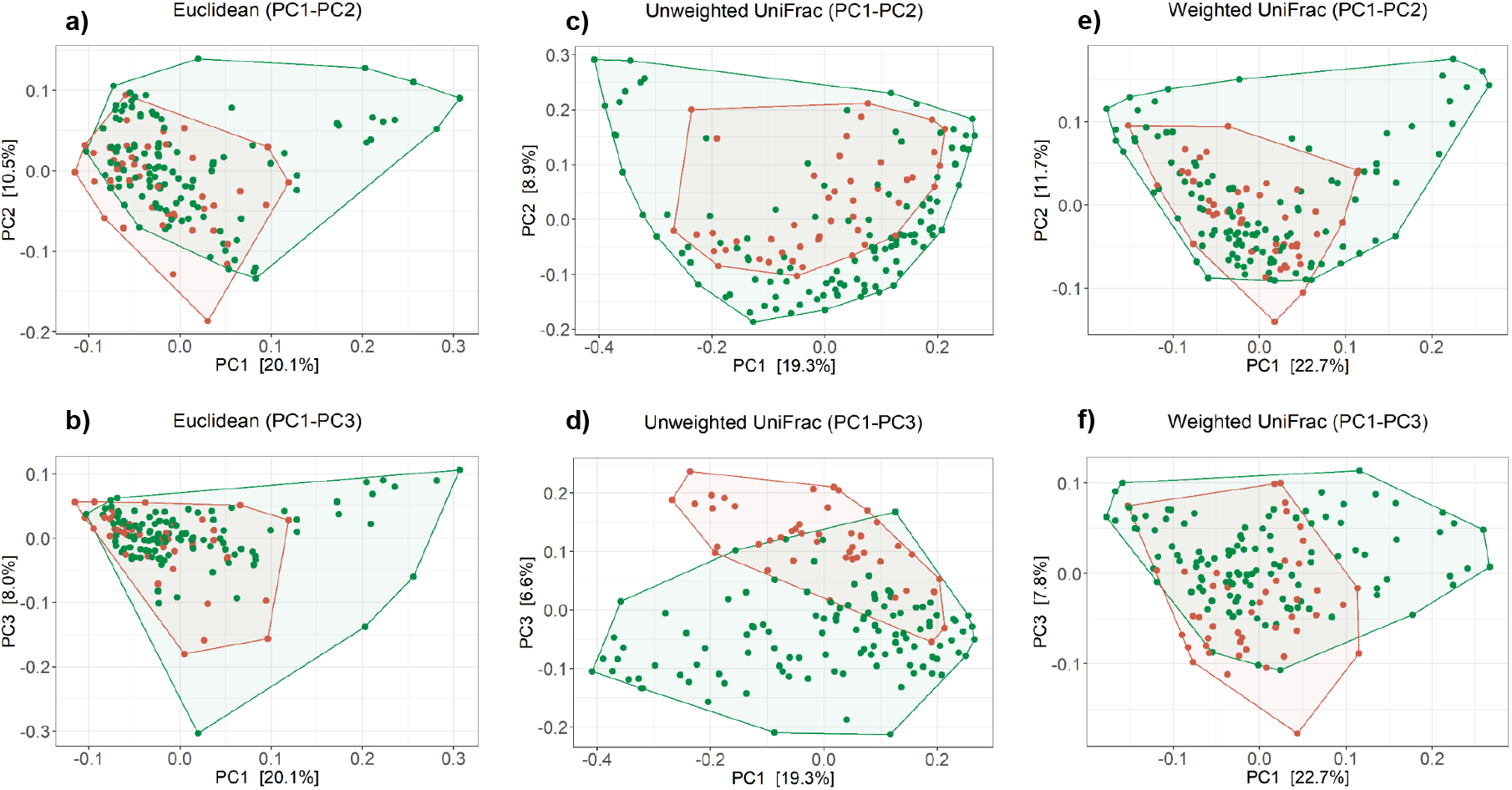
Composition of bacterial community in the gut of mouse lemurs differs between habitats. Principal-coordinate plots of Euclidean (a) PC1-PC2, (b) PC1-PC3; Unweighted UniFrac (c) PC1-PC2, (d) PC1-PC3); Weighted UniFrac (e) PC1-PC2, (f) PC1-PC3) metrics in mouse lemurs. Dots and connecting polygons represent bacterial communities in gut of mouse lemurs trapped in Andranovao (green) or Miarintsoa (red).

### Relative abundance of major phyla and ASVs differ between habitats

The habitat influenced the relative abundance of taxa at various taxonomic levels. At the higher phylum level, five phyla, namely Verrucomicrobia, Fusobacteria, Spirochaetes, Lentisphaerae and Tenericutes, were detected as being differentially abundant (p<0.01) between Andranovao and Miarintsoa individuals. All observed phyla showed a decrease in the relative abundance in Miarintsoa compared with Andranovao (Figure 4). Among the detected phyla, Verrucomicrobia was found to be the top ranked phylum according to the Random forest classifier. At the ASV level, we identified 48 ASVs that differed significantly (p<0.01) in abundance in mouse lemurs, with 33 OTUs (68.7%) revealing a lower and 15 OTUs (31.2%) revealing a higher mean abundance in Miarintsoa compared with Andranovao individuals (Figure 5, Supplementary Table S3).

**Figure 4.**
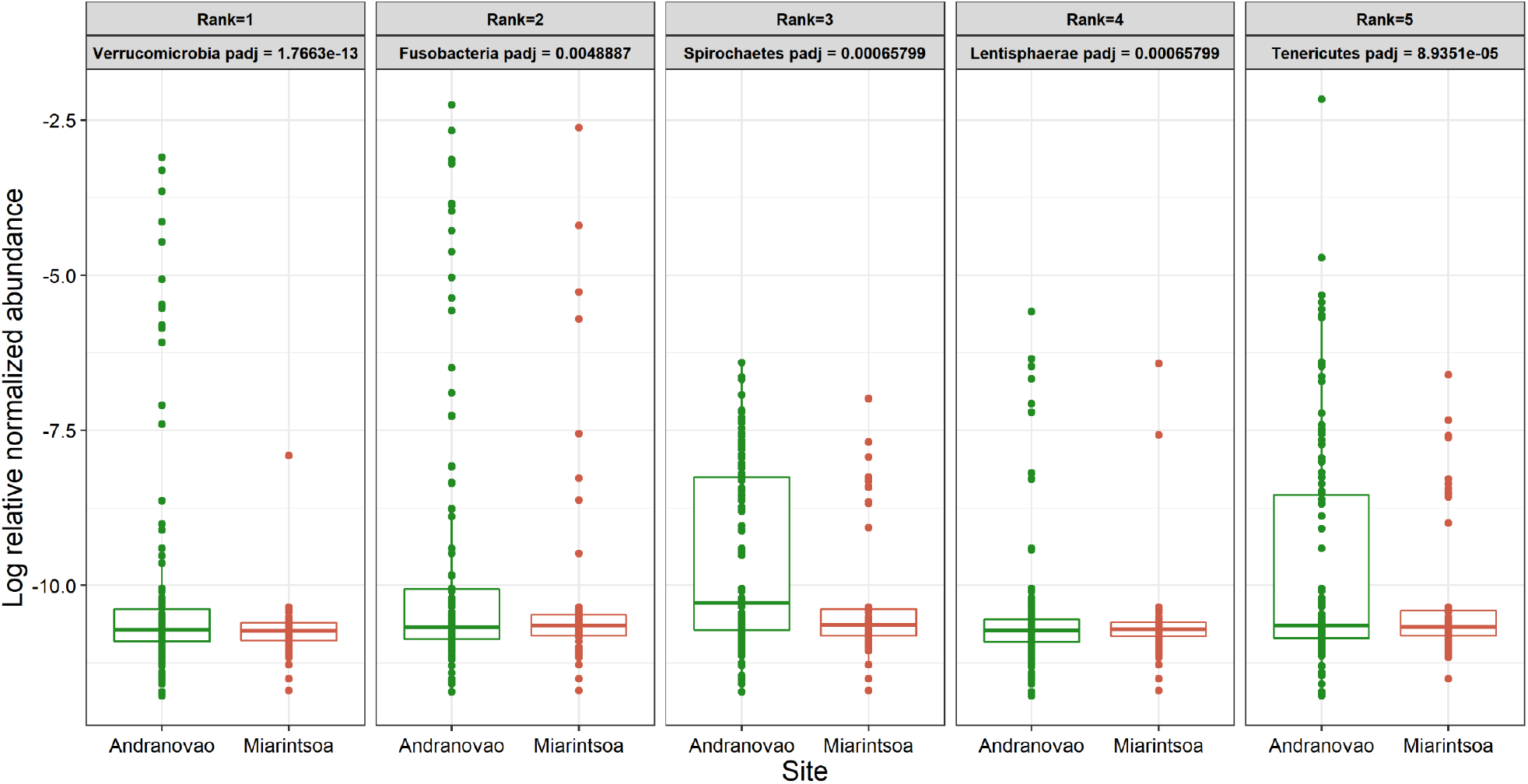
Relative abundance of major bacterial phyla in gut of mouse lemurs differs between habitats. Box plots indicate the effect of study site (Andranovao in green; Miarintsoa in red) on the relative abundance of major phyla in the gut microbiomes of mouse lemurs (all p<0.01). Phyla are arranged according to the assigned rank based on the importance of differentially abundant phyla (see methods for details).

**Figure 5.**
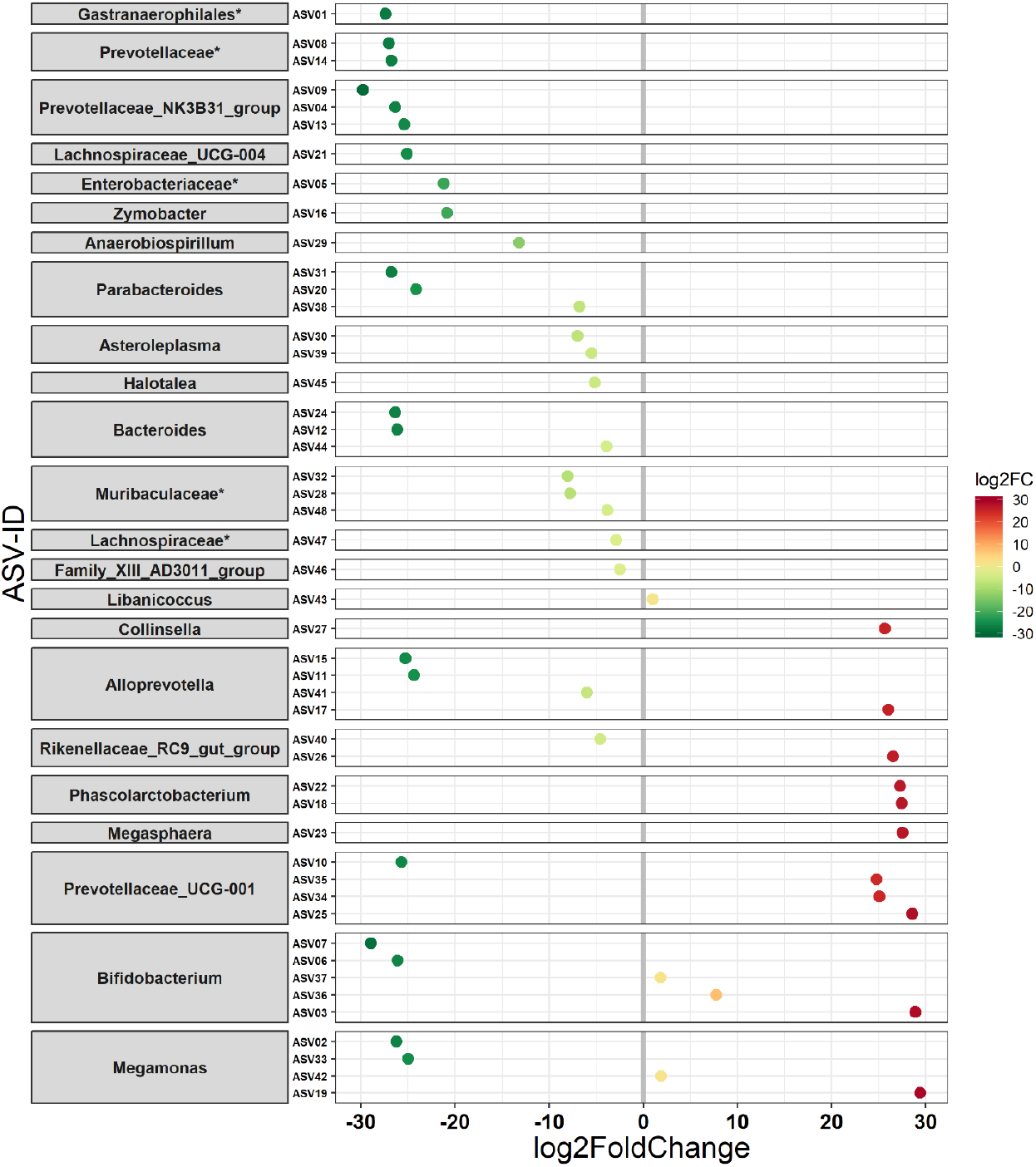
Differential abundance of ASVs in mouse lemurs inhabiting two different habitats. ASVs (48 ASVs) that differ in their mean abundance in relation to the habitat of mouse lemurs. Values indicate a log two-fold (log2FC) decrease (33 ASVs) or increase (15 ASVs) in Miarintsoa individuals. ASVs are arranged according to increasing values of log2FC. The highest possible taxonomic assignment (maximal to the genus level) is shown for each ASV; *includes unclassified ASV/s at genus level.

ASVs that showed a lower abundance in mouse lemurs from Miarintsoa predominantly belonged to the families Prevotellaceae (n=9 ASV), Bacteroidaceae (n=3 ASV), Tannerellaceae (n=3 ASV), Muribaculaceae (n=3 ASV) and genera *Prevotellaceae_NK3B31_group* (n=3 ASV), *Parabacteroides* (n=3 ASV) and *Bacteroides* (n=3 ASV) (Figure 5, Supplementary Table S3). ASVs that showed a higher abundance in Miarintsoa individuals mainly belonged to the families Prevotellaceae (n=4 ASV), Bifidobacteriaceae (n=3 ASV), Veillonellaceae (n=3 ASV) and genera *Prevotellaceae_UCG-001* (n=3 ASV) and *Bifidobacterium* (n=3 ASV) (Figure 5, Supplementary Table S3).

### Predicted metagenomes and higher functional pathways differ between lemurs from two sites

We applied a PERMANOVA-based model approach to investigate whether differences in the predicted KEGG orthologs (KOs) occurred in relation to site (habitat disturbance) or sex. In the model, site (R^2^=0.012, p=0.039) had a significant effect on the *gower* distances, but not sex (R^2^=0.003, p=0.941).

Four predicted higher functional pathways (KEGG) were identified that differed significantly in their abundance between Andranovao and Miarintsoa individuals (p≤0.05) (Supplementary Figure 1). At the higher level, all the identified pathways were related to “metabolism”. The identified pathway that showed severely lower abundance (25 fold) in Miarintsoa individuals belonged to “Metabolism of terpenoids and polyketides”. The metabolism pathway, “Xenobiotics biodegradation and metabolism”, also showed a decrease in individuals from Miarintsoa, whereas the abundance of two other metabolic pathways (“Carbohydrate metabolism”, “Metabolism of other amino acids”) increased in Miarintsoa compared with Andranovao individuals (Supplementary Figure 1).

## Discussion

Despite major conservation efforts, Madagascar’s unique ecosystems are declining at a vast speed, threatening the home of many endemic lemur species. Whereas most lemur species are highly sensitive to anthropogenic change, others, including our focus species, which is one of the smallest Malagasy primates, namely the grey-brown mouse lemur, (*M. griseorufus*), seem at the first glance to be more resilient, since they can also inhibit human-modified habitats (Hending, 2021). However, whether such resilience also applies to its inner ecosystem, the gut microbiome, remains unknown. Nevertheless, increasing numbers of studies emphasize the negative impacts of habitat modification on the homeostasis of the intestinal microbiome, which is key to host health (Amato et al., 2013; Barelli et al., 2015, 2020; Hayakawa et al., 2018; Trosvik et al., 2018). Reduction in overall gut microbial diversity can result in ‘dysbiosis’, i.e. a depletion in bacteria essential for host health and an increase in pathogenic bacteria, both of which can negatively affect the health of grey-brown mouse lemurs (Duvallet et al., 2017).

Our comparison of the gut microbiomes of mouse lemurs inhabiting two different habitats, namely Andranovao situated in Tsimanampetsotsa National Park and the highly disturbed habitat Miarintsoa surrounded by human settlements, revealed that anthropogenic disturbance was indeed associated with the disruption of the homeostasis of the gut microbiome. This was reflected by a decrease in microbiome alpha and beta diversity and an alteration in microbial community composition in Miarintsoa. Moreover, we observed a decrease in beneficial bacterial taxa and associated shifts in the predicted metabolic functions of the microbiome of the grey-brown mouse lemurs living in the disturbed habitat. Even though Miarintsoa receives more rain per year (∼100mm difference) than Andranovao (about 40km distance), which might provide a higher abundance and diversity of fruit plants and should give rise to higher gut microbial diversity, we observed a reduction in gut microbial diversity suggesting that the influence of anthropogenic disturbance is greater than that of better food availability attributable to the wetter climate.

Habitat modifications can stimulate many factors impacting the host microbial community and its functions: (1) diet-associated factors driven by a change in vegetation arising from (micro)climatic, rainfall or land-use shifts, (2) altered home range sizes caused by habitat disturbance and changes in food items and their distribution, and (3) proximity to humans, livestock and other domestic animals (Wasimuddin et al., 2017; Fackelmann et al., 2021). In Andranovao, the natural vegetation consists of largely intact dry spiny forest and spiny thicket, whereas in Miarintsoa, which usually receives more rainfall, the vegetation is dominated by agricultural fields and pasture with small and scattered forest remnants used intensively for wood collection (Yedidya Ratovonamana et al., 2011, 2013). Such disturbances in forests over time might constrain the movement of mouse lemurs, restrict its ranging pattern (Bohr et al., 2011; Steffens and Rakotondranary, 2017) and impact its natural diet composition. Reduction and modification in diet diversity has also been suggested to cause the decreases seen in the gut microbial diversity of other non-human primates (Amato et al., 2013; Barelli et al., 2015, 2020; Hayakawa et al., 2018; Trosvik et al., 2018). Similarly, in association with increasing disturbance, an increase in the activity period of mouse lemurs in order to meet their energy demands has been noted in disturbed forests (Fish 2014). All such behavioural and physiological changes can disturb the diversity and composition of the gut microbiome of mouse lemurs but cannot be disentangled in this study.

Consistent with the decrease in overall gut microbial diversity in mouse lemurs from Miarintsoa, five phyla (Verrucomicrobia, Fusobacteria, Spirochaetes, Lentisphaerae and Tenericutes) had a lower relative abundance in the disturbed habitat compared with individuals from the National Park. All of these phyla are common in various non-human primate guts (Clayton et al., 2018). The presence of Verrucomicrobia and Fusobacteria in human guts has proven beneficial for host health (Ghosh and Pramanik, 2021). Interestingly, higher proportions of phyla Verrucomicrobia and Lentisphaerae have been positively correlated with better sleep and high cognitive abilities (Anderson et al., 2017). Thus, the reduced abundance of these two phyla might explain the increased activity in lemurs living in fragmented forests (Fish, 2014). Furthermore, the lower abundance of these phyla makes the host susceptible to other gut disturbances (Kalkeri et al., 2021). Spirochaetes are normally present in the gut microbiomes of humans and other mammals and play an important role in the digestion of plant polysaccharides (Thingholm et al., 2021). Moreover, in baboons, social interactions are crucial for the transmission of gut bacteria, specifically anaerobic and non-spore-forming bacteria such as Spirochaetes, between members of the same social group (Tung et al., 2015). One can speculate that the observed reduced abundances of Spirochaetes in Miarintsoa individuals is attributable to the limited contact that they have with other co-specifics because of habitat disturbance. Little is known so far about the functional role of members of the Tenericutes phylum; however, they are commonly present in the gut of non-human primates, humans, fishes and insects (Clayton et al., 2018; Wang et al., 2020b). Nevertheless, a deeper understanding of the cause-and-effect relationships for phyla abundance requires further investigations.

Among the ASVs that differed significantly between the Miarintsoa and Andranovao individuals, most ASVs were lower (68%) in mouse lemurs from Miarintsoa. At the family level, these decreasing ASVs predominantly belonged to Prevotellaceae followed by Bacteroidaceae, Tannerellaceae and Muribaculaceae and, at the genera level, decreasing ASVs were assigned to the *Prevotellaceae_NK3B31_group, Parabacteroides* and *Bacteroides*. The family Prevotellaceae is associated in humans and chimpanzees with their carbohydrate-rich diets; this is also true for grey-brown mouse lemurs whose diet is primarily based on gum and fruits (Moeller et al., 2012). Interestingly, the *Prevotellaceae_NK3B31_group* has been suggested to provide protection against invading pathogens and to reduce diarrhoea in piglets (Wang et al., 2020a). Thus, the reduced relative abundance of members of the Prevotellaceae family in mouse lemurs from the disturbed habitat Miarintsoa might be a result of their ingesting a diet less rich in carbohydrates but richer in insects, which ultimately might make mouse lemurs more susceptible to invading pathogens. Bacteroidaceae and its genus *Bacteroides*, and *Parabacteroides* (a genus of Tannerellaceae) are generally considered as beneficial bacteria with numerous benefits to host health (Wexler and Goodman, 2017). They are major producers of many short chain fatty acids in the gut and play an important role in gut homeostasis and immune regulation (Venegas et al., 2019; Wang et al., 2019). A decrease in the abundance of Bacteroidaceae/*Bacteroides* and *Parabacteroides* has been demonstrated in various intestinal disorders (Forbes et al., 2016).

Mouse lemurs represent an important model in aging research because of their high phenotypic plasticity and similarity with human aging (Hozer et al., 2019). Interestingly, members of the Muribaculaceae family are often reported to be associated with aging and longevity (Sibai et al., 2020). Thus, a decrease in the relative abundance of members of the Muribaculaceae in Miarintsoa might be a sign of accelerated aging and/or disturbed biological rhythms in anthropogenically disturbed landscapes. However, we have also noted an increase of a smaller number of ASVs (31%) mainly belonging to the Prevotellaceae family (genus *Prevotellaceae_UCG-001*), Bifidobacteriaceae family (genus *Bifidobacterium*) and members of the Veillonellaceae family in the gut of mouse lemurs from Miarintsoa. Genera of the Prevotellaceae family are well known for their competitive interactions and thus a decrease in the abundance of some of their members and increase of others is not surprising (Lei et al., 2018). However, the genus *Bifidobacterium* is usually considered as beneficial for host health and therefore an increase in the abundance of its members in mouse lemurs trapped in a disturbed habitat is surprising (Sun et al., 2020). Members of the Veillonellaceae family are often found in association with gut inflammation (Ling et al., 2016) and hence are more abundant in patients with Inflammatory Bowel Disease, fibrosis and other disease (Forbes et al., 2016; Lee et al., 2020; Salliss et al., 2021). Overall, the decrease in the abundance of beneficial bacterial taxa together with an increase in taxa associated with a diseased state implies a loss of homeostasis in the gut microbiomes of mouse lemurs from Miarintsoa; this might adversely affect the health of lemurs and potentially make them more susceptible to diseases.

In accordance with the changes in the microbial community composition, namely the decrease in bacterial diversity and in the beneficial taxa in mouse lemurs captured in the disturbed habitat, we detected shifts in microbial metabolic functions. All predicted microbial functional pathways differing between lemurs of Miarintsoa and Andranovao sites belonged to metabolism at a higher level. The major pathway that showed severely lower abundance (25 fold) in Miarintsoa individuals was assigned to “Metabolism of terpenoids and polyketides”. Terpenoids and polyketides are secondary metabolites from plants and trees and a reduced abundance of functions related to their metabolism might constrain the digestive abilities of the hosts (Grassotti et al., 2021). Similarly, the pathway related to “Xenobiotics biodegradation and metabolism” was also noted to be lower in Miarintsoa individuals.

Because of the crucial services performed by the microbiome with regard to host health, such negative alterations of the gut microbiome of lemurs living in anthropogenically disturbed habitats might impact the health of even a phenotypically resistant species, rendering them susceptible to diseases and ultimately impacting their survival in Madagascar’s shrinking biodiversity. Investigation of the gut microbiome represents a non-invasive and simple method for detecting early signs of declining wildlife health and is easy to implement as a diagnostic tool in primate conservation.

## Acknowledgements

The study was carried out under the collaboration between Madagascar National Parks, the Department of Animal Biology, Department of Plant Biology and Ecology (Antananarivo University, Madagascar) and the Department of Animal Ecology and Conservation (Hamburg University, Germany). We are grateful to all field assistants for their constant help during our fieldwork. We thank Kerstin Wilhelm for excellent technical assistance. This research was funded by SuLaMa/BMBF (FKZ 01LL0914) and the German Science Foundation (DFG) and is part of the DFG Priority Program SPP 1596/2 Ecology and species barriers in emerging infectious diseases (GA 342/19-1, DR 772/8-1, SO 428/9-1).

## Data Availability Statement

The individual gut bacterial 16S rRNA gene sequences are available under NCBI SRA accession ID SRP217185.

## Contributions

S.S., J.U.G. and W. conceived the study. J.U.G., J.R. and Y.R.R. provided host data. W. carried out laboratory experiments, bioinformatics and statistical analyses. H.M. contributed to draft scripts. W. wrote the MS. All authors commented on and approved the final version of the MS.

## Supplementary Figure and Tables

**Supplementary Figure 1.**
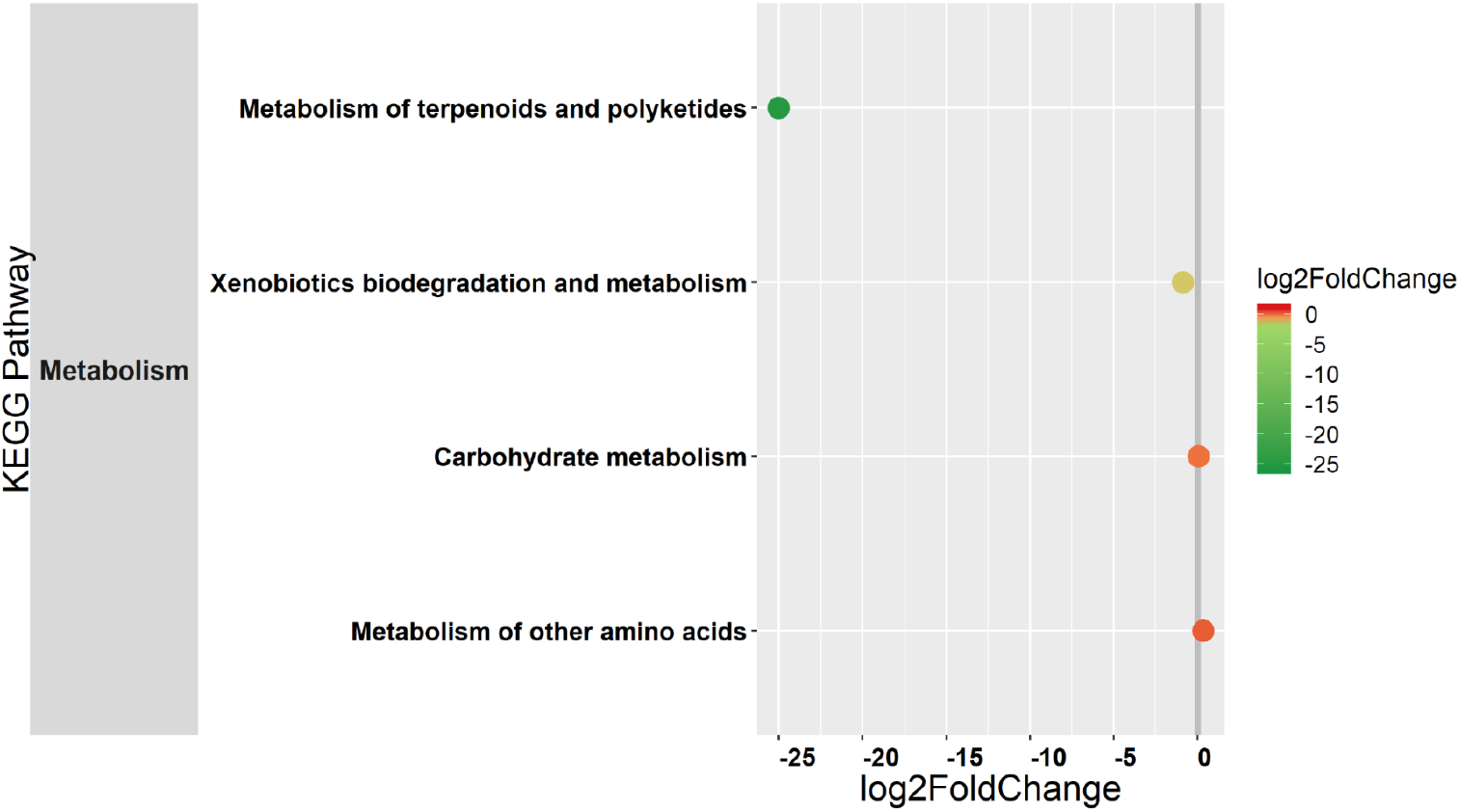
Differential abundance of predicted major functional pathways in relation to the habitat of mouse lemurs. Differences in the mean abundance of major functional pathways (by using KEGG classification) between Andranovao and Miarintsoa individuals (Wald tests, p≤0.05). The values indicate a log two-fold decrease or increase in Miarintsoa individuals. Functional pathways are arranged according to increasing values of log two-fold change.

**Supplementary Table 1**. Summary of the best model (GLM) predicting (a) observed species, (b) Fisher diversity and (c) Shannon diversity according to habitat and sex of the mouse lemurs. Significant values (<0.05) are shown in bold.

**Supplementary Table 2**. Summary of the PERMANOVA models based on (a) Euclidean, (b) Unweighted UniFrac and (c) Weighted UniFrac metric according to habitat and sex of the mouse lemurs. Significant values (<0.05) are shown in bold.

**Supplementary Table 3**. Nomenclature and abundances of 48 ASVs differing in their mean abundance in relation to the habitat of the mouse lemurs. Only highly significant ASVs (after FDR correction, ≤0.01) are shown; they are arranged according to increasing log two-fold change (log2FC) values. In the column log2FC, ASVs that showed a lower abundance (i.e. mean read coverage) in mouse lemurs captured in the disturbed site Miarintsoa than in those in Andranovao (located in the Tsimanampetsotsa National Park) are marked in green (68.7%); those that increased in abundance in Miarintsoa individuals are marked in red (31.2%).

